# Discrimination of SARS-CoV-2 Omicron sub-lineages BA.1 and BA.2 using a high-resolution melting-based assay: A pilot study

**DOI:** 10.1101/2022.04.11.487970

**Authors:** Akira Aoki, Hirokazu Adachi, Yoko Mori, Miyabi Ito, Katsuhiko Sato, Kenji Okuda, Toru Sakakibara, Yoshinori Okamoto, Hideto Jinno

**Affiliations:** Faculty of Pharmacy, Meijo University, 150 Yagotoyama, Tempaku-ku, Nagoya 468-8503, Japan; Aichi Prefectural Institute of Public Health, 7-6 Nagare, Tsuji-machi, Kita-ku, Nagoya 462-8576, Japan; Chita Health Center, 88-2 Arako-ushiro, Yawata-aza, Chita, Aichi 478-0001, Japan

**Keywords:** SARS-CoV-2, Receptor-binding domain, Omicron variant, BA.1, BA.2, High-resolution melting

## Abstract

The Omicron variant of severe acute respiratory syndrome coronavirus 2 (SARS-CoV-2) has spread worldwide. As of March 2022, Omicron variant BA.2 is rapidly replacing variant BA.1. As variant BA.2 may cause more severe disease than variant BA.1, variant BA.2 requires continuous monitoring. The current study aimed to develop a novel high-resolution melting (HRM) assay for variants BA.1 and BA.2 and to determine the sensitivity and specificity of our method using clinical samples. Here, we focused on the mutational spectra at three regions in the spike receptor-binding domain (RBD; R408, G446/L452, and S477/T478) for the variant-selective HRM analysis. Each variant was identified based on the mutational spectra as follows: no mutations (Alpha variant); L452R and T478K (Delta variant); G446S and S477N/T478K (Omicron variant BA.1); and R408S and S477N/T478K (Omicron variant BA.2). Upon analysis of mutation-coding RNA fragments, the melting peaks of the wild-type fragments were distinct from those of the mutant fragments. The sensitivity and specificity of this method were determined as 100% and more than 97.5%, respectively, based on 128 clinical samples (40 Alpha, 40 Delta, 40 Omicron variants BA.1/BA.1.1, and 8 Omicron BA.2). These results suggest that this HRM-based assay is a promising screening method for monitoring the transmission of Omicron variants BA.1 and BA.2.

## INTRODUCTION

The severe acute respiratory syndrome coronavirus 2 (SARS-CoV-2) has spread worldwide, and novel coronavirus disease 2019 (COVID-19) cases continue to rise rapidly. Multiple SARS-CoV-2 variants have been detected (1, 2), and some spike receptor-binding domain (RBD) mutations, such as N501Y and L452R, can increase viral infectivity and cause immune escape. The Alpha (B.1.1.7) and Delta (B.1.617.2) variants, respectively, harboring N501Y and L452R, greatly contributed to the COVID-19 pandemic. In late September 2021, the novel Omicron variant (B.1.1.529), emerged from South Africa and spread worldwide rapidly (3, 4). This variant has many spike mutations (5), and the vaccine breakthrough infection rate of the Omicron variant is reportedly higher than that of other variants (6, 7). Thus, we should strive to prevent the spread of Omicron infection, even though the vaccination program is proceeding.

Recent studies have reported that Omicron sub-lineages, such as BA.1, BA.2, and BA.3, have emerged in several countries (8, 9). Since late 2021, the BA.1 variant (including BA.1.1, which harbors R346K) has become the main Omicron sub-lineage worldwide. In early 2022, the BA.2 variant showed an increasing prevalence in some countries, such as Denmark and the United Kingdom (10, 11), while the BA.3 variant is unlikely to spread. Because the spike deletion at 69-70 was recognized as a signature mutation of the Omicron variant, it was used to detect the Omicron variant. However, the BA.2 variant lacks this deletion, meaning that it goes undetected (12). The BA.2 variant was then called “Stealth Omicron.” A preliminary report demonstrated that the reproduction number of BA.2 was 1.4-fold higher than that of BA.1, and BA.2 was more pathogenic in hamsters than BA.1 (13). Therefore, the BA.2 variant may outcompete BA.1 in the near future, and BA.2 should be monitored with caution.

Next-generation sequencing (NGS) has been used to diagnose SARS-CoV-2 variants (14); however, this process requires several days to determine the genome sequence. Rapid screening tests can help diagnose any SARS-CoV-2 variant within a day. Our previous reports showed that a screening assay using high-resolution melting (HRM) analysis, a post-PCR technique for genotyping based on the melting behavior of amplicons, could identify the Omicron (B.1.1.519) and Delta variants (15, 16). We successfully developed an HRM-based assay to identify Omicron B.1.1.529 and Delta mutants. At two RBD regions, namely G446/L452 and S477/T478, the assay could identify the mutational spectra; G446S/L452 and S477N/T478K indicated Omicron variants, while G446/L452R and S477/T478K indicated Delta variants (Table 1). Although the S477N/T478K RBD substitutions are common mutations among Omicron sub-lineages, BA.2 possesses some unique RBD mutations, including R408S without the G446S substitution.

**Table 1.** Receptor-binding domain (RBD) amino acid substitutions in Alpha, Delta, and Omicron variants.

| WHO label | Pangolin | RBD amino acid substitutions |
| --- | --- | --- |
| Alpha | B.1.1.7 | N501Y |
| Delta | B.1.617.2 | **L452R, T478K** |
| Omicron | B.1.1.529/BA.1 | G339D, S371L, S373P, S375F, K417N, N440K, **G446S**,<br>**S477N, T478K**, E484A, Q493R, G496S, Q498R,<br>N501Y, Y505H |
| Omicron | BA.2 | G339D, S371F, S373P, S375F, T376A, D405N, **R408S**,<br>K417N, N440K, **S477N, T478K**, E484A, Q493R,<br>Q498R, N501Y, Y505H |
| Omicron | BA.3 | G339D, S371F, S373P, S375F, D405N, K417N, N440K,<br>**G446S, S477N, T478K**, E484A, Q493R, Q498R,<br>N501Y, Y505H |

Substitutions detected by this high-resolution melting assay are marked as bold.

In this study, we developed an HRM-based assay to distinguish BA.1 (including BA.1.1) and BA.2 at another RBD site, R408. Moreover, we validated this HRM-based assay at three RBD regions, R408, G446/L452, and S477/T478, using 128 clinical samples (n = 40 Alpha variant, n = 40 Delta variant, n = 40 Omicron variant BA.1/BA.1.1, n = 8 Omicron variant BA.2).

## MATERIALS AND METHODS

### Ethics statement

This project was approved by the Research Ethics Committee of Meijo University (Approval number: 2020-17-2) and Aichi Prefectural Institute of Public Health (Approval number: 20E-4) and was carried out according to the Infectious Diseases Control Law of Japan.

### Preparation of standard RNA fragments: *In vitro* T7 transcription

The SARS-CoV-2 sequence was obtained from NCBI (GenBank ID: MN908947), the GISAID database (www.gisaid.org), and the Pango nomenclature system (https://cov-lineages.org/lineages.html). Five RBD DNA fragments (wild type, R408S mutant, BA.1 variant mutant, BA.2 variant mutant, Delta variant mutant; 600-1000 bp in length) with a 5′ T7 upstream promoter sequence were obtained from Eurofins Genomics K.K. (Tokyo, Japan). *In vitro* T7 transcription was performed as described previously (16). The synthesized single-stranded RNA fragments were used as reverse transcriptase (RT)-PCR amplification templates.

### RT-PCR amplification: First PCR

RT-PCR was performed in a single closed tube using a one-step RT-PCR kit (One Step PrimeScript III RT-qPCR Mix, with UNG; Takara Bio Inc., Kusatsu, Japan) in accordance with the manufacturer’s instructions. The primer pairs used were as follows: R408 outer forward 5′-TGCTTGGAACAGGAAGAGAA-3′ and R408 outer reverse 5′-AACGCAGCCTGTAAAATCATC-3′; G466-T478 outer forward 5′-TTACAGGCTGCGTTATAG-3′ and G466-T478 outer reverse 5′-ACAAACAGTTGCTGGTGCAT-3′ (Fig. 1). Each DNA fragment was observed as a single, correctly-sized band as follows: R408, 247 bp; G466-T478, 290 bp. RT-PCR amplification was performed as previously described (16). After amplification, the reaction mixture was diluted 10,000-fold with water and used as a template for the second PCR and HRM analyses.

**Fig. 1.**
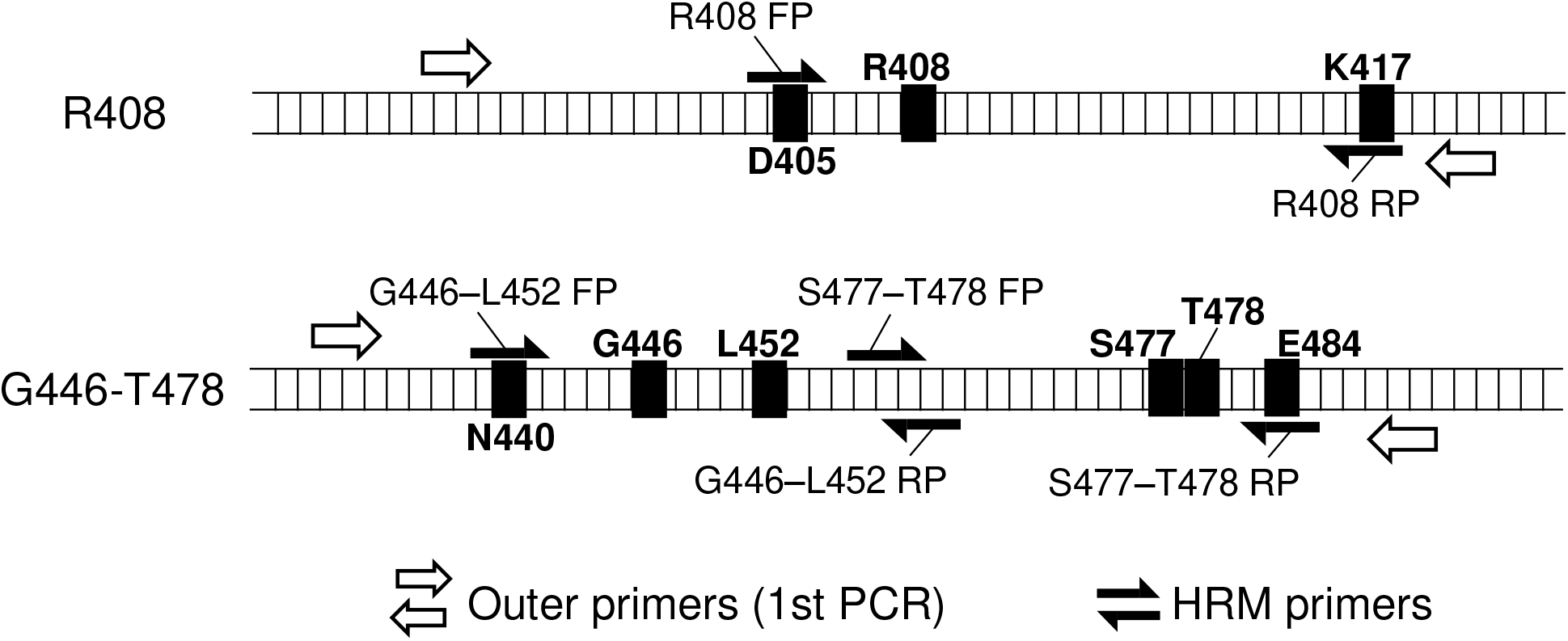
Schematic map of primer annealing sites for RT-PCR and high-resolution melting analyses.

### HRM analysis: Second PCR

HRM was performed using an HRM reagent (MeltDoctor HRM Master Mix; Thermo Fisher Scientific, Waltham, MA, USA) according to the manufacturer’s instructions. The second primer pair was as follows: R408 forward 5′-TGCAGATTCATTTGTAATTAGAGGTGATGAAG-3′ and R408 reverse 5′-CTTTCCAGTTTGCCCTGGAG-3′; G446-L452 forward 5′-GGCTGCGTTATAGCTTGGAATTCTAACAATCTT-3′ and G446-L452 reverse 5′-TCAAAAGGTTTGAGATTAGACTTCC-3′; S477-T478 forward 5′- TTGTTTAGGAAGTCTAATCTCAAACC-3′ and S477-T478 reverse 5′-AAGTAACAATTAAAACCTTCAACACCATTACAAGG-3′. Each DNA fragment was observed as a single, correctly-sized band as follows: R408, 64 bp; G466-L452, 104 bp; S477-T478, 107 bp. As shown in Fig. 1, the R408 forward primer design was based on the D405 coding sequence, R408 reverse primer design was based on the K417 coding sequence, G446–L452 forward primer design was based on the N440 coding sequence, and S477–T478 reverse primer design was based on the E484 coding sequence to avoid the potential influences of D405, K417, N440, and E484 mutations, respectively. Briefly, each reaction mixture (20 μL) contained 2 μL of the diluted RT-PCR mixture, 400 nmol/L of each primer, and 1 × master mix. All reactions were performed in duplicate using a real-time PCR system (LightCycler 96 System; F. Hoff Mann-La Roche Ltd., Basel, Switzerland). PCR amplification was performed as previously described (16). HRM curves were analyzed using Gene Scanning Software, version 1.1.0.1320 (F. Hoffmann-La Roche Ltd.) under default settings. The normalized melting curve and melting peaks (−dF/dT) were acquired by setting the pre-melt and post-melt fluorescence at 100% and 0%, respectively.

### Clinical samples

From July 2021 to January 2022, 128 nasopharyngeal swabs or saliva samples were collected from patients suspected of having COVID-19 and those detected with COVID-19 by Aichi Prefectural Institute of Public Health. The Ct values in the clinical quantitative PCR test for these samples ranged from 15 to 35. Viral RNA was extracted from clinical samples using spin columns (QIAamp Viral RNA Mini QIAcube Kit; Qiagen GmbH, Hilden, Germany) and an automated nucleic acid purification system (QIAcube Connect; Qiagen). Each eluent was reverse-transcribed, PCR-amplified, and subjected to library preparation using the QIAseq FX library kit (Qiagen), according to the protocol established by the National Institute of Infectious Diseases, Tokyo, Japan (17, 18). Whole-genome sequencing was performed using a next-generation sequencer (MiSeq system; Illumina Inc., San Diego, CA, USA). The obtained NGS reads were mapped to the SARS-CoV-2 Wuhan-Hu-1 reference genome sequence (29.9 kb single-strand RNA; GenBank ID: MN908947). For lineage assignment, Pangolin version 3.1.20, with pangoLEARN version 2022–02-28 was used.

## RESULTS

### HRM analysis of standard RNA fragments

Although the Omicron sub-lineages BA.1 and BA.2 were found to be unique RBD substitutions, there was no BA.3-specific substitution in the RBD (Table 1). At present, there are far fewer new cases of the BA.3 variant than of either BA.1 or BA.2. Hence, the main aim of this assay is to discriminate between BA.1 and BA.2. In this study, we developed an HRM-based assay for the R408 site, where BA.2 has an R408S substitution but not BA.1. The normalized melting curves and melting peaks for R408, R408S, and BA.2 RBD are shown in Fig. 2. The R408S RBD plot was different from the R408 RBD plot. In addition, the BA.2 RBD plot was in good agreement with the R408S plot. Then, BA.2 RBD was analyzed with the HRM assay at G446/L452R and S477/T478, which we developed previously (15). The HRM assay correctly classified BA.2 as a G446/L452 and S477N/T478K variant (Figs. S1 and S2). These results suggest that our HRM assay can identify BA.2 and discriminate between BA.1 and BA.2 in the three RBD regions.

**Fig. 2.**
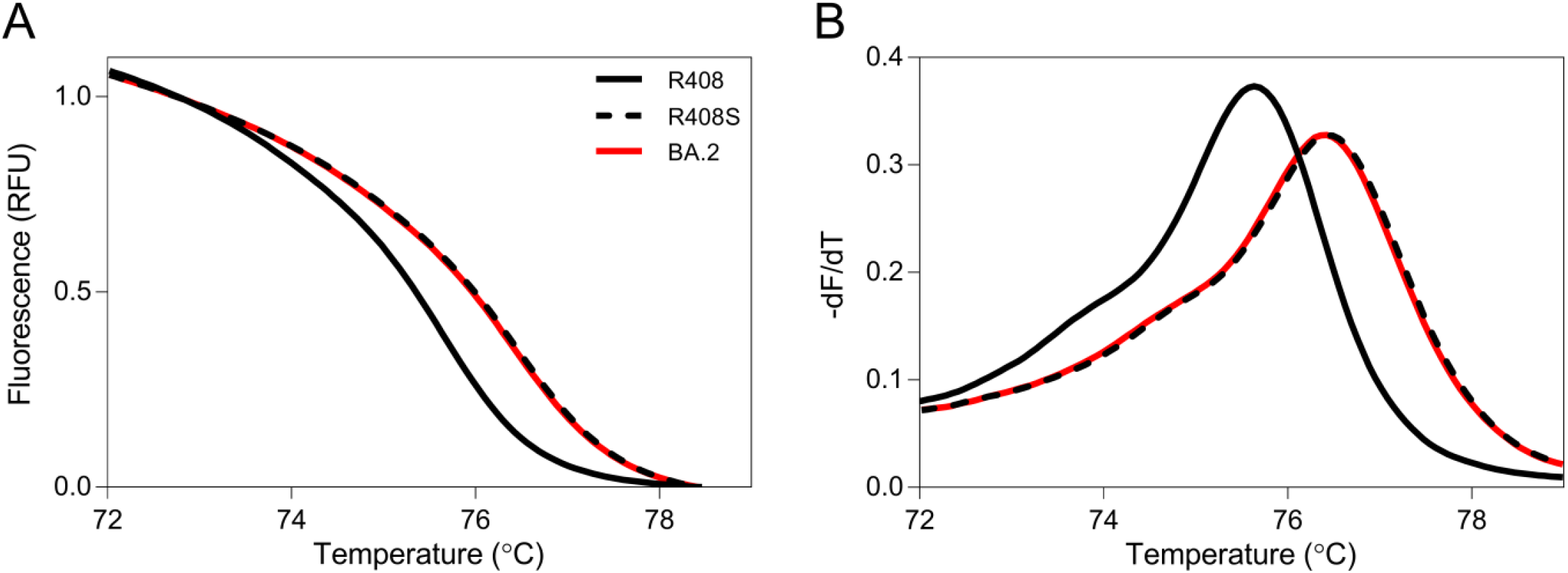
Normalized melting curves and melting peaks of positive control RNAs for the R408 site. Normalized melting curve plots (A) and melting peak plots (B) for the R408 site were acquired using standard fragments of the R408 receptor-binding domain (RBD; solid black line), R408S RBD (dashed black line), and BA.2 RBD (solid red line).

### HRM analysis of clinical samples

Finally, we confirmed the applicability of this HRM analysis to clinical samples. The Alpha variant was found to have no RBD substitutions at the R408/T478 sites (Table 1). Thus, this HRM-based assay identified a clinical sample containing the Alpha variant as the wild-type. After whole-genome sequencing by NGS, 128 clinical samples (n = 40 Alpha, n = 40 Delta, n = 40 BA.1/BA.1.1, n = 8 BA.2 variants) were randomly selected (Table 2). Clinical samples were analyzed using HRM analysis, together with positive control RNAs. After HRM analysis, Gene Scanning Software automatically classified samples into wild-type or mutant strains based on HRM curves. The results of the HRM analysis of three regions, R408, G446/L452, and S477/T478, are shown in Table 3, and the melting peak plots of representative samples are shown in Fig. 3. At the R408 site, all eight BA.2 samples were correctly classified as the R408S mutant, and all others were classified as wild-type. At the G446/L452 site, 40 BA.1 variant samples were classified as the G446S mutant, and 40 Alpha samples, 40 Delta samples, and 8 BA.2 samples were correctly classified as the wild-type. At the S477/T478 site, all BA.1 and BA.2 samples were correctly classified as S477N/T478K mutants. One Alpha sample and one Delta sample plot disagreed with each positive control plot (Fig. 3). Therefore, we calculated the sensitivity and specificity of the current HRM-based assay based on these results. The sensitivity and specificity of the assay for the three regions were 100% and over 97.5%, respectively (Table 4).

**Table 2.** Pangolin of 128 clinical samples by whole-genome sequencing.

| Alpha |  | Delta |  | Omicron |  |
| --- | --- | --- | --- | --- | --- |
| Pangolin | No. of samples | Pangolin | No. of samples | Pangolin | No. of samples |
| B.1.1.7 | 40 | AY.29 | 37 | BA.1 | 6 |
|  |  | AY.29.1 | 1 | BA.1.1 | 34 |
|  |  | AY.46 | 1 | BA.2 | 8 |
|  |  | AY.92 | 1 |  |  |

**Table 3.**
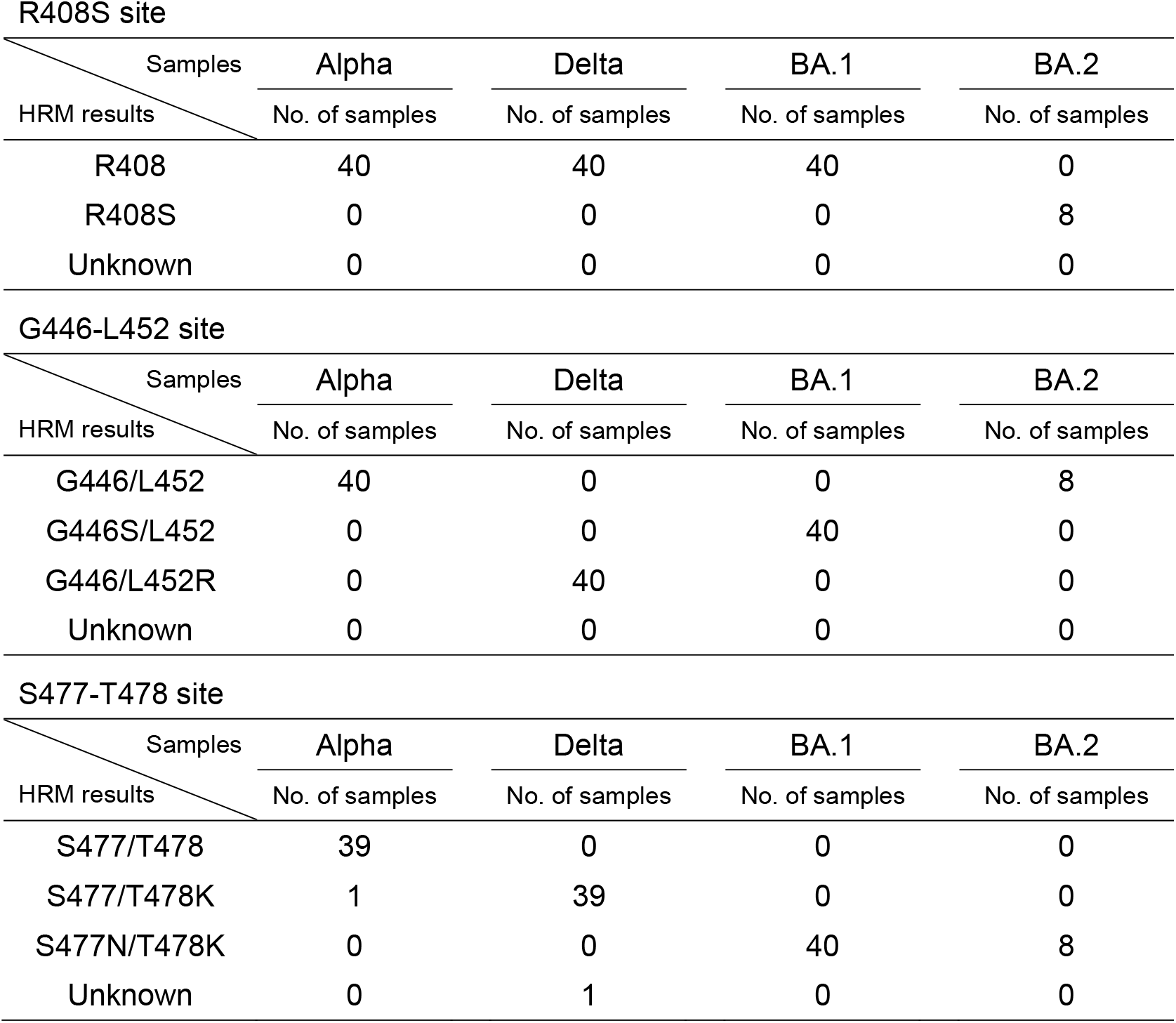
Results of high-resolution melting analysis at R408, G446/L452, and S477/T478 sites.

| Samples<br>HRM results | Alpha | Delta | BA.1 | BA.2 |
| --- | --- | --- | --- | --- |
|  | No. of samples | No. of samples | No. of samples | No. of samples |
| R408 | 40 | 40 | 40 | 0 |
| R408S | 0 | 0 | 0 | 8 |
| Unknown | 0 | 0 | 0 | 0 |

G446-L452 site
| Samples<br>HRM results | Alpha | Delta | BA.1 | BA.2 |
| --- | --- | --- | --- | --- |
|  | No. of samples | No. of samples | No. of samples | No. of samples |
| G446/L452 | 40 | 0 | 0 | 8 |
| G446S/L452 | 0 | 0 | 40 | 0 |
| G446/L452R | 0 | 40 | 0 | 0 |
| Unknown | 0 | 0 | 0 | 0 |

S477-T478 site
| Samples<br>HRM results | Alpha | Delta | BA.1 | BA.2 |
| --- | --- | --- | --- | --- |
|  | No. of samples | No. of samples | No. of samples | No. of samples |
| S477/T478 | 39 | 0 | 0 | 0 |
| S477/T478K | 1 | 39 | 0 | 0 |
| S477N/T478K | 0 | 0 | 40 | 8 |
| Unknown | 0 | 1 | 0 | 0 |

**Table 4.** Sensitivity and specificity of the high-resolution melting assay in comparison with next-generation sequencing.

|  | R408S detection | G446S detection | S477N/T478K detection |
| --- | --- | --- | --- |
| Sensitivity<sup>a</sup> | 100% (8 / 8) | 100% (40 / 40) | 100% (48 / 48) |
| Specificity<sup>b</sup> | 100% (120 / 120) | 100% (88 / 88) | 97.5% (78 / 80) |

<sup>a</sup>Sensitivity: True positive number/(True positive number+False Positive number)x100
<sup>b</sup>Specificity: False negative number/(False negative number+True negative number)x100

**Fig. 3.**
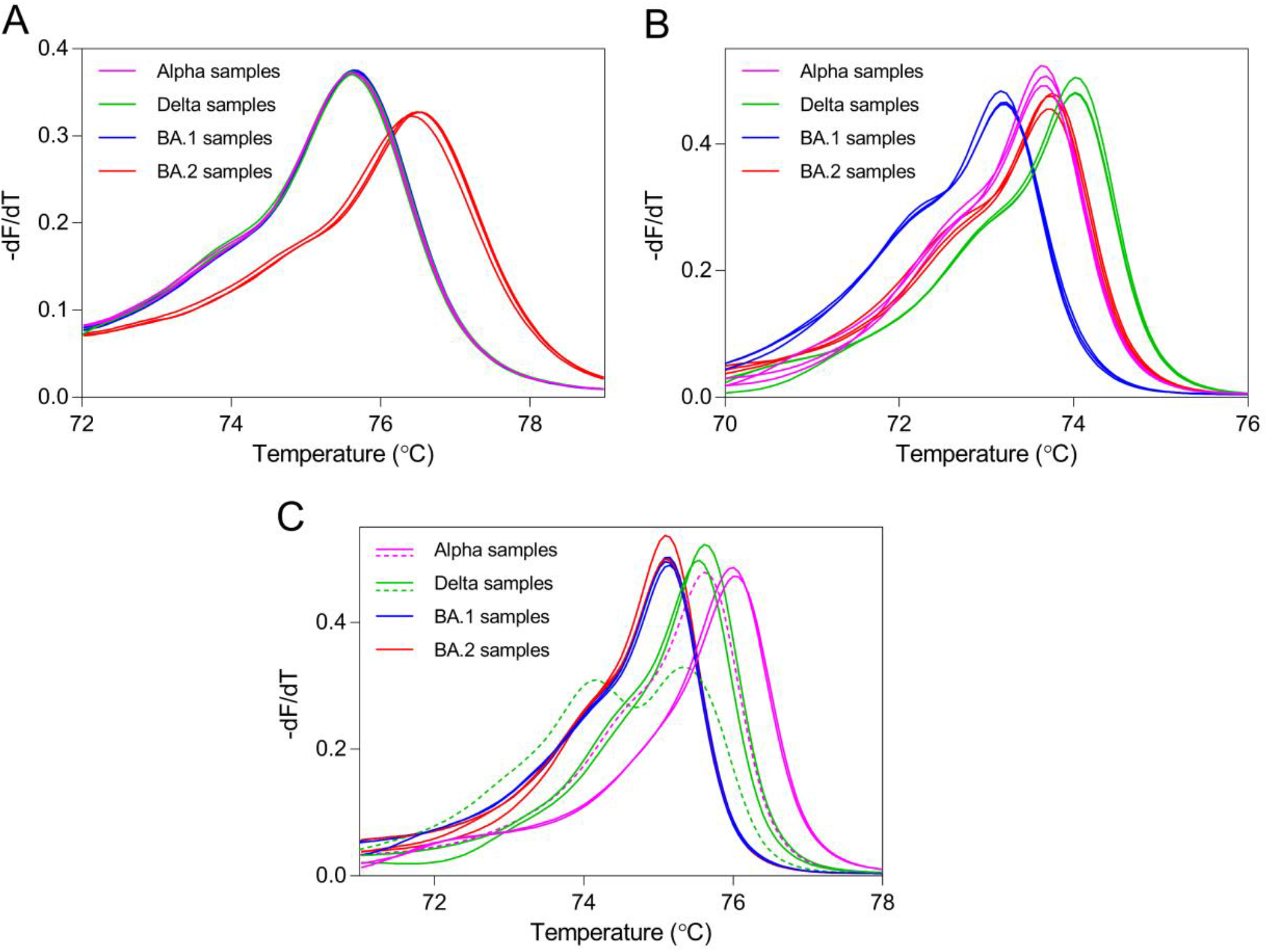
Melting peaks of clinical samples for three receptor-binding domain regions. Melting peak plots for the R408 (A), G446/L452 (B), and S477/T478 sites were acquired using clinical samples with three Alpha variants (pink line), three Delta variants (green line), three BA.1/BA.1.1 variants (blue line), and three BA.2 variants (red line). Solid lines indicate true positive and true negative samples. Dashed lines indicate false negative samples.

## Discussion

This study is the first report developing the screening test for discriminating Omicron variants BA.1 and BA.2 using HRM assay. This HRM-based assay had high sensitivity and specificity for BA.2 diagnosis. Additionally, the Delta variant was identifiable using this assay. New cases of infection with the Delta variant, such as the AY.29 sub-lineage (19), remain detectable in some countries, including Japan. The Delta variant increases the risk of hospitalization than the Omicron variant (20). Therefore, rigorous global monitoring of both Omicron and Delta variants is warranted.

RBD is the essential region in which mutations lead to increase infectivity and reduce antibody affinity. All SARS-CoV-2 variant of concern has characteristic RBD substitutions. In contrast, many mutations, excluding the RBD region, are not common among similar lineages, as the genomic sequence of SARS-CoV-2 continues to mutate over time. Direct detection of characteristic RBD mutations contributes to avoiding misdiagnosis of each variant. Therefore, we have developed HRM assays to detect SARS-CoV-2 RBD mutations (15, 16). Moreover, HRM analysis can detect unknown mutations unlike TaqMan probe assays. If a post-Omicron variant emerges, we can quickly develop an HRM-based assay to detect the RBD mutation in the new variant.

The TaqMan probe assay is typically used for single-nucleotide polymorphism detection. Multiple TaqMan probe kits for the detection of SARS-CoV-2 mutations are commercially available worldwide. As the Omicron RBD is highly mutated, the reactivity of the available specific probes may be affected, which is likely to increase the risk of false negatives. This HRM-based assay without a specific probe can be constructed more easily than a TaqMan probe assay. Thus, HRM analysis is useful in identifying highly mutated variants. However, the HRM assay may not be able to identify samples that contain low virus copies. Our previous study showed that when used with samples containing >10^7^ copies/mL, the HRM assay could detect SARS-CoV-2 mutations (21). In this study, the two-step nested PCR improved the detection limit of the HRM assay, and low-copy virus samples could be identified, such as those having Ct values of 35. Moradzad et al. (22) reported that a one-step HRM assay can detect SARS-CoV-2 mutations with relatively high sensitivity and specificity, similar to the TaqMan probe assay. However, they used high-copy number viral RNAs (10 ng RNA in a tube) for HRM analysis. Additional work is required to improve the detection limit of our HRM-based assay using a one-step single-tube PCR assay.

This study needs to be interpreted in the context of its limitations. First, this study was performed on clinical samples from a limited area of Japan. Further studies are needed to confirm the utility of this HRM-based assay using a larger size of samples from various regions. Second, the low-copy virus samples with a Ct value less than 35 may more frequently result in a false positive or false negative. The limit of detection should be determined using low-copy virus samples. Third, our assay cannot discriminate between Omicron variants BA.1 and BA.3 because these sub-lineages possess the same mutational spectra at R408, G446S/L452, and S477N/T478K (Table 1). Even if the variant BA.3 replaces BA.1 and BA.2, we can identify BA.3 by determining the D405N mutation in combination with others.

In conclusion, we developed a novel assay to identify the main Omicron sub-lineages BA.1 and BA.2, using HRM analysis. As this HRM-based genotyping assay does not require sequence-specific probes, unlike the TaqMan probe assay, it is easy to perform and is cost-effective. Our results suggest that the current HRM-based assay is a powerful high-throughput tool for determining the SARS-CoV-2 Omicron sub-lineages. Since this assay was verified using limited clinical samples, further studies using diverse samples are needed to validate this HRM-based assay among the various institutions.

## ACKNOWLEDGMENTS

The authors would like to thank the staff of the Laboratory of Virology, Department of Microbiology and Medical Zoology, Aichi Prefectural Institute of Public Health, who performed COVID-19 PCR testing, RNA purification from clinical samples, and NGS analysis.

All authors participated in the interpretation of the results. A.A. designed and conducted HRM analysis and wrote the original draft of the manuscript. H.A. and M.I. conducted NGS analysis and edited the manuscript. Y.M. performed sequence analysis. Y.O. planned the study and edited the manuscript. K.S., K.O., T.S. and H.J. planned the study and supervised the project. All the authors critically reviewed and approved the final version of the manuscript.

This work was supported in part by the DAIKO FOUNDATION and Meijo University Research Project for Countermeasures Against COVID-19.

## FIGURE LEGENDS

**Fig. S1.**
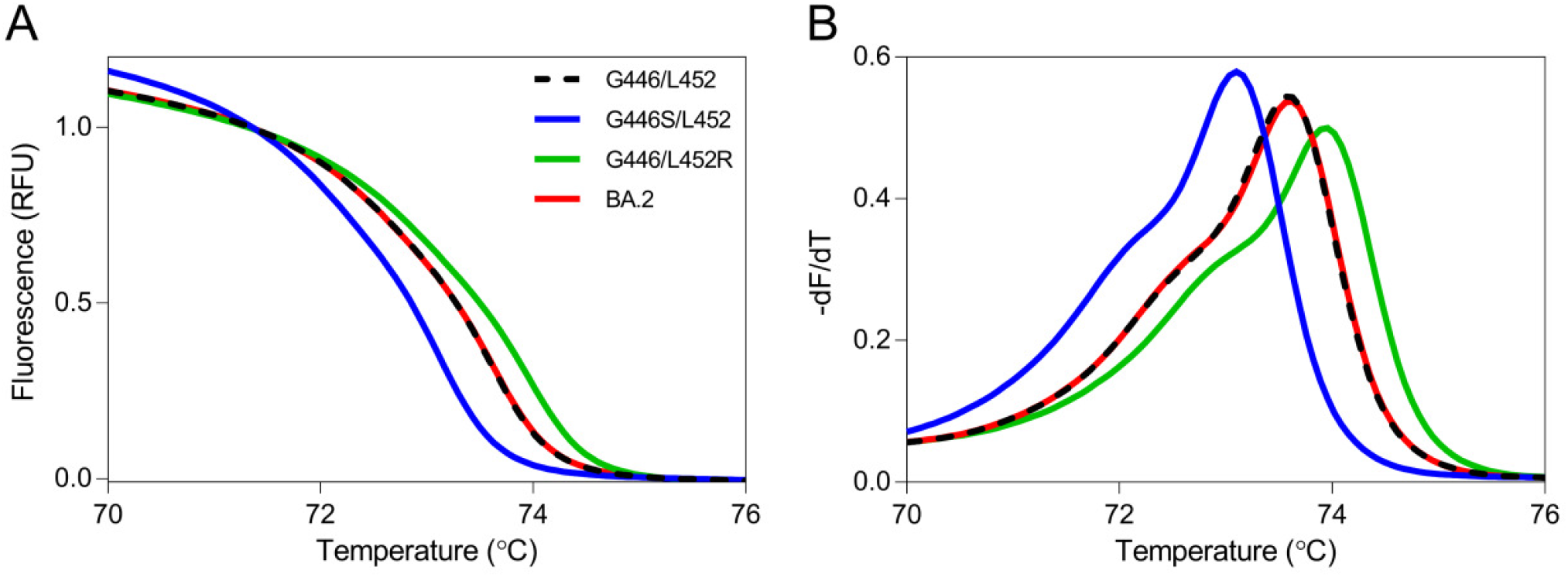
Normalized melting curves and melting peaks of positive control RNAs for the R446/L452 site. Normalized melting curve plots (A) and melting peak plots (B) for the R446/L452 site were acquired using standard fragments of the G446/L452 RBD (dashed black line), G446S/L452 RBD (solid blue line), G446/L452R RBD (solid green line), and BA.2 RBD (solid red line).

**Fig. S2.**
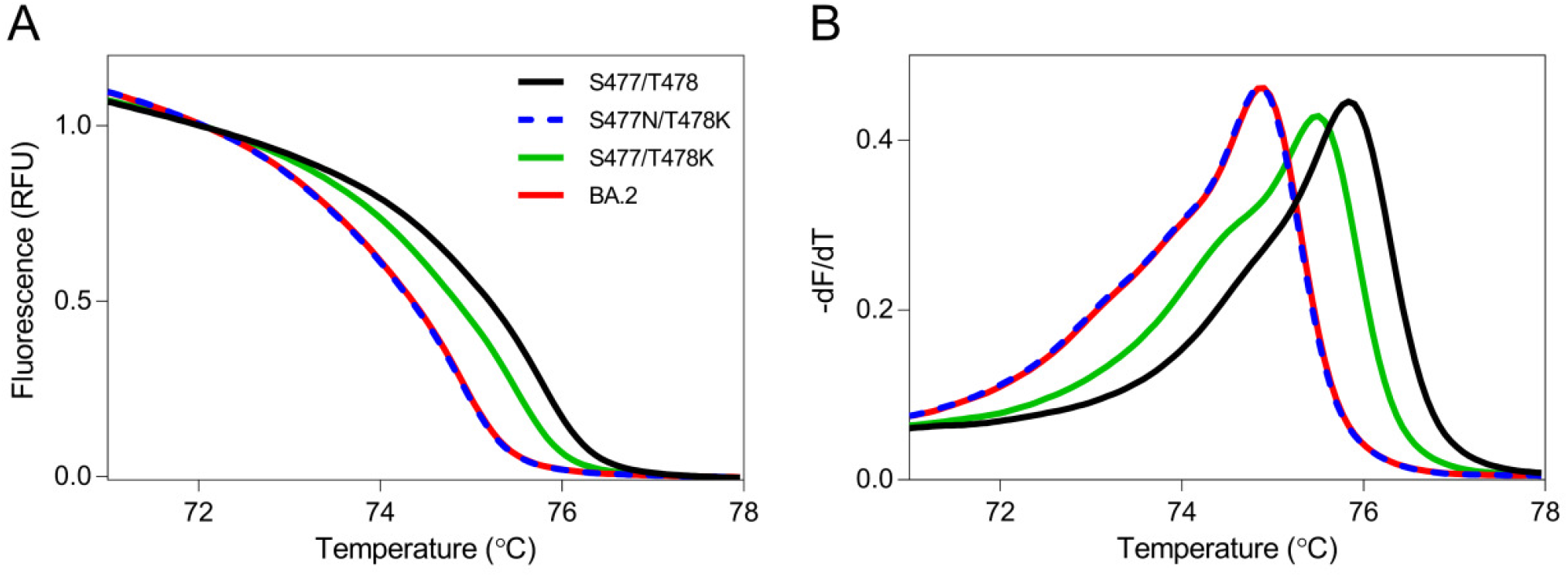
Normalized melting curves and melting peaks of positive control RNAs for the S477/T478 site. Normalized melting curve plots (A) and melting peak plots (B) for the S477/T478 site were acquired using standard fragments of the S477/T478 RBD (solid black line), S477/T478 RBD (dashed blue line), S477/T478 RBD (solid green line), and BA.2 RBD (solid red line).

